# Intestinal uptake regulates T cell responses to dietary antigens

**DOI:** 10.64898/2026.09.01.748748

**Authors:** Ya-Ting Chang, Henry H. Le, Andrew Morrison, Ryan Kong, Ell Schulman, Tsui-Wen Chou, Jamie E. Blum

## Abstract

Dietary proteins induce antigen-specific immune responses, leading to oral tolerance or food allergies. While mouse models of allergic sensitization rely on antigen context and adjuvant signaling, exposure to purified proteins is usually sufficient for tolerance development in studies with soluble model antigens. In previous work, we discovered that the maize protein zein induces robust antigen-specific intestinal Tregs following normal chow exposure, but unexpectedly does not elicit a T cell response when delivered as a purified protein. Here, we investigated the mechanisms underlying these differential T cell responses, considering both biochemical properties of zein and its food matrix, as well as intestinal antigen processing. We focused on three zein preparations that induced different frequencies of antigen-specific intestinal T cells. While *in vitro* studies showed comparable presentation by dendritic cells, *in vivo* studies revealed stark differences in intestinal uptake. We identified differences in protein solubility and active sampling mechanisms as cooperative drivers of intestinal zein uptake. Different zein preparations could be sampled by either goblet cells, a route previously described for the model antigen OVA, or M cells, a route hypothesized for dietary proteins and established for transport of pathogenic bacteria. These findings also extend to peanuts, where the allergen Ara h 1 was selectively transported by goblet or M cells depending on food preparation, establishing examples of naturally occurring dietary M cell ligands. Finally, solubilizing zein increased intestinal uptake, and correspondingly feeding a diet with solubilized zein led to more intestinal zein-specific T cells compared to feeding the insoluble form. Overall, these findings suggest that intestinal uptake of dietary antigens is a regulated and context-dependent determinant of antigen-specific T cell induction, with implications for development of tolerance-restoring immunotherapies.

## Introduction

Dietary antigens are actively sampled across the intestinal epithelium and screened by the intestinal immune system, where encountered antigens largely result in oral tolerance, the process of immunologically annotating dietary components as safe^1,2^. However, although most interactions are benign, dietary antigens can also elicit potent allergic responses. Antigen-specific immune responses are established through the activation and differentiation of naïve T cells, where development into a Treg leads to oral tolerance^3^. Conversely, development into Th2 and Tfh effector cells coordinates allergic sensitization^4,5^. Identifying factors that guide T cell activation and differentiation to regulatory or effector fates is thus critical to decoding immune interpretation of food antigens.

Allergy development is context-dependent in mice, with the conditions of an antigen exposure strongly influencing whether sensitization occurs^6^. For example, co-administered adjuvants (e.g., cholera toxin, SEB) are often required to induce allergic sensitization, where in most cases, exposure to purified protein is not sufficient to elicit allergy^7–10^. Additionally, cooking and processing can also change protein allergenicity^11,12^. In contrast, how the context of antigen exposure and form of an antigen influence oral tolerance is less well understood. Treg induction can be enhanced by a select set of bile acids, as well as dietary components such as fiber and arginine^13–15^. However, while these interventions enhance the Treg response to protein feeding, tolerance is still induced in mice fed purified protein alone. Thus, these studies suggest that oral administration of protein alone is sufficient to induce tolerance. However, since prior studies of immune tolerance have focused on a narrow set of soluble model antigens, it is unclear how food matrix components, physical antigen properties, or other features of antigen exposure may broadly influence the development of oral tolerance.

In recent work, we determined that the maize protein αZein is an immunodominant dietary antigen that induces a sizeable intestine-resident Treg population (>2% of pTregs) in C57BL/6 mice^16^. Surprisingly, although feeding a zein-containing chow diet elicited a robust intestinal T cell response, a diet with purified zein induced little zein-specific T cell development^16^. Thus, this example highlights a case where a dietary antigen that elicits a robust immune response within a food matrix is largely immunologically silent when delivered as a purified protein (**Fig 1A**). Motivated by this finding, in this study, we use zein to evaluate antigen sufficiency and context dependency for immune tolerance toward dietary antigens. While we originally anticipated identifying maize-derived molecules that activate innate immunity, our data instead revealed that the route of intestinal uptake may influence the extent of antigen-specific T cell development and tolerance induction.

**Figure 1:**
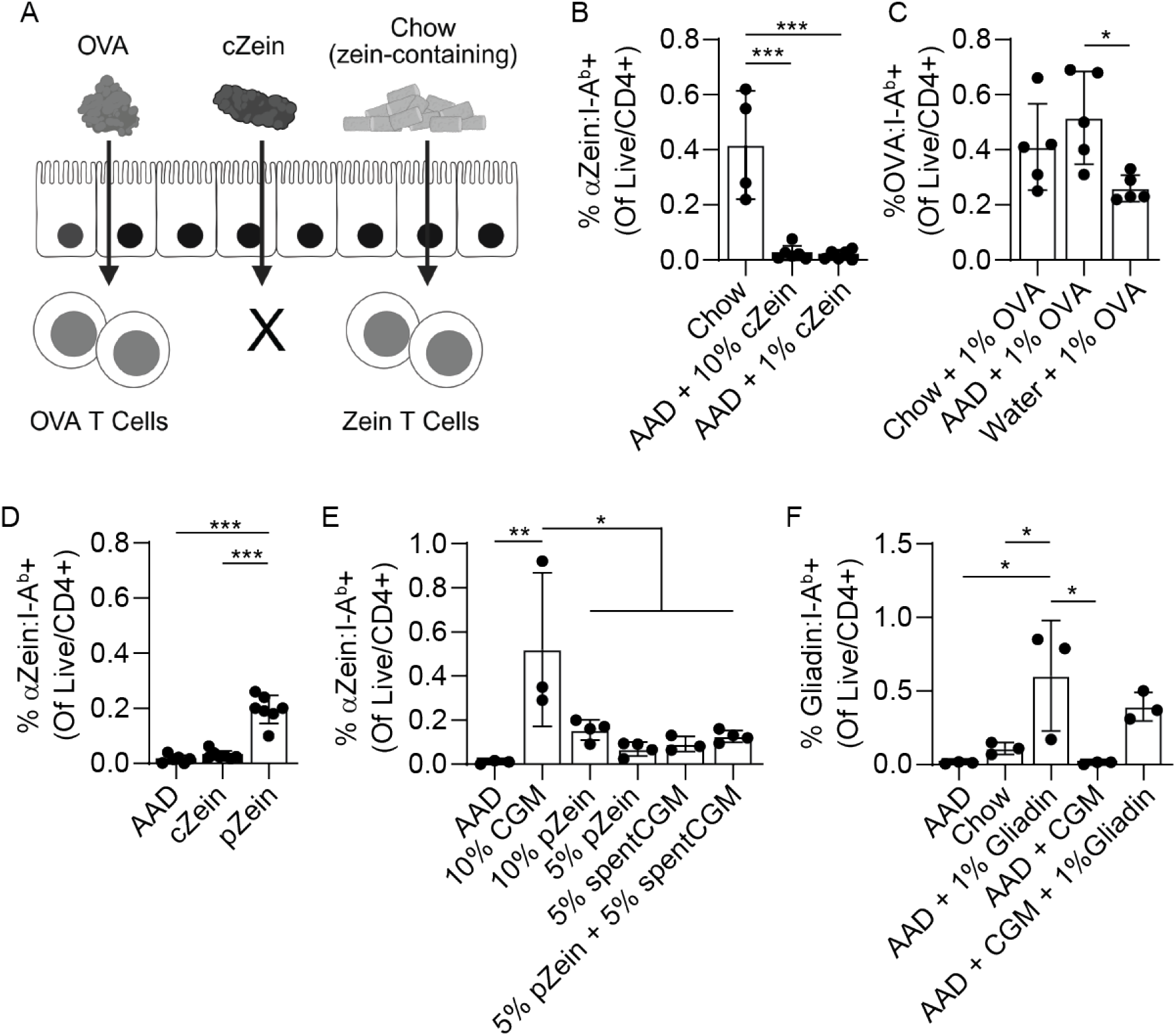
Zein-containing diets show different T cell induction capacities. **(A)** Intestinal antigen-specific T cells can be selectively induced in response to purified protein or whole food feeding. **(B)** Intestinal zein-specific T cells in mice fed chow, AAD+10% cZein, or AAD+1% cZein (N=4-6/group). **(C)** Intestinal OVA-specific T cells in mice fed chow+1% OVA, AAD+1% OVA, or water+1% OVA (N=5/group). **(D)** Intestinal zein-specific T cells in mice fed AAD, AAD+10% cZein, or AAD+10% p Zein (N=6-7/group). **(E)** Intestinal zein-specific T cells in mice fed AAD, AAD+10% CGM, AAD+10% pZein, AAD+5% pZein, AAD+5% spentCGM, AAD+5% pZein + 5% spentCGM (N=3-4/group). **(F)** Intestinal gliadin-specific T cells in mice fed AAD, chow, AAD+1% gliadin, AAD+CGM, AAD+CGM+1% gliadin (N=3/group). * P < 0.05, ** P < 0.01, *** P < 0.001. Error bars represent standard deviation.

## Results

### Dietary or protein context is critical for αZein T cell induction

Confirming our prior work, we found that commercially acquired purified zein (henceforth, cZein) was insufficient to induce the development of zein-specific T cells *in vivo* (**Fig 1B, Fig S1**). This unexpected finding seemingly contradicts the established paradigm with the model antigen ovalbumin (OVA), which induces a T cell response when delivered as purified protein^3,17,18^. We confirmed the sufficiency of purified OVA for T cell induction across delivery in drinking water or incorporation into different diets (**Fig 1C**). In fact, with adoptively transferred OVA-specific T cells (OT-II cells), an amino acid defined (AAD) diet increased abundance of OT-II cells relative to chow diet, consistent with a prior report, and likely reflecting a lack of competing intestinal antigens (**Fig S2A**)^17^. We discovered that feeding a diet containing purified zein from a second vendor was sufficient for intestinal zein-specific T cell development (**Fig 1D**). This second zein is extracted from corn gluten meal (CGM), a protein-rich by-product of cornstarch and corn oil production, and from this point forward will be referred to as processed zein (pZein). By mass, CGM contains approximately 50-70% zein proteins, with the remaining component being water-insoluble fiber and hydrophobic metabolites^19^. Compared to pZein, CGM more robustly induced zein-specific intestinal T cells (**Fig 1E**). The immunogenicity of CGM could reflect a change in the zein protein, such as processing-induced chemical modifications; an adjuvant effect of the non-zein components; or a physical matrix effect. To evaluate these factors, we measured whether reconstituting pZein with “spentCGM” (the remaining components of CGM following pZein extraction) could restore zein-specific T cell induction. Recombining pZein with spentCGM did not increase zein-specific T cell numbers, suggesting that a co-occurring adjuvant molecule is not sufficient to explain differences in zein-specific T cell development between CGM and pZein (**Fig 1E, Fig S2B-C**). In all cases, the induced zein-specific T cells adopted a Treg phenotype, suggesting that food context alters T cell abundance but does not change the resulting phenotypic response (**Fig S2D**). Lastly, we sought to determine whether the apparent food context dependency was specific to zein or shared among other proteins with similar biochemical properties (i.e., ethanol soluble proteins). To address this, we compared the T cell response to gliadin delivered alone (AAD + gliadin), in chow (naturally wheat-containing), or alongside CGM (AAD + gliadin + CGM). Similar to OVA, purified gliadin was sufficient to induce gliadin-specific T cells (**Fig 1F**). Additionally, co-administration of CGM and gliadin did not enhance the gliadin-specific T cell response, confirming that CGM does not behave as a classical adjuvant. Taken together, these results highlight that various zein preparations result in different T cell responses in an adjuvant-independent manner, suggesting that protein context beyond the adjuvant effect plays a role in immune system processing.

### Protein composition and particle properties between preparations

We next aimed to determine properties of the zein preparations that underlie potential interactions with the immune system and differential T cell induction capacity. Since our data argue against a classical adjuvant effect, variation in zein-specific T cell induction could instead reflect differences in the zein protein itself, such as composition or processing-induced modifications, or in physical properties such as size or surface charge. Zein protein in corn is a family of over 30 unique zein isoforms with minor differences in amino acid sequence. Notably, only some isoforms contain the 11 amino acid T cell epitope. Thus, compositional differences could explain differences in corresponding cognate T cell induction. However, LCMS-based proteomics revealed that the top zein isoforms were shared across each zein preparation, with no loss of the antigenic epitope in any preparation (**Fig 2A, Table S1**). Additionally, in an *in vitro* system where primary APCs are used to stimulate T cell lines heterologously expressing a zein-specific TCR, cZein, pZein, and CGM induced comparable T cell activation, further revealing that the zein preparations harbor unaltered epitopes which equally stimulate APC antigen presentation and T cell activation (**Fig 2B**). Protein features such as particle size and charge can alter how proteins are perceived by the immune system, however particle size and zeta potential did not identify obvious differences (**Fig 2C-D**)^20,21^. Upon dissolving the various mass-normalized zein sources in simulated intestinal fluid (SIF), we noted that CGM and pZein produce more soluble zein than cZein (**Fig 2E**). Zein solubility can be controlled by surface features and aggregation state, independent of size and charge^22^. Thus, we hypothesized that differences in particle architecture may change solubility. Supporting this possibility, SEM revealed considerable morphological differences between pZein and cZein. Notably, cZein appears as smooth and rounded particles, while pZein particles appear with angular edges and thin sheet-like structures, which potentially provide more solvent-accessible surface thus increasing solubility (**Fig 2F**). Further, glutelin proteins were enriched in pZein compared to cZein (**Fig 2A**). Loss of glutelins in cZein may reflect additional wash steps in the extraction process that both remove contaminants and simultaneously change particle architecture, thereby modifying solubility.

**Figure 2:**
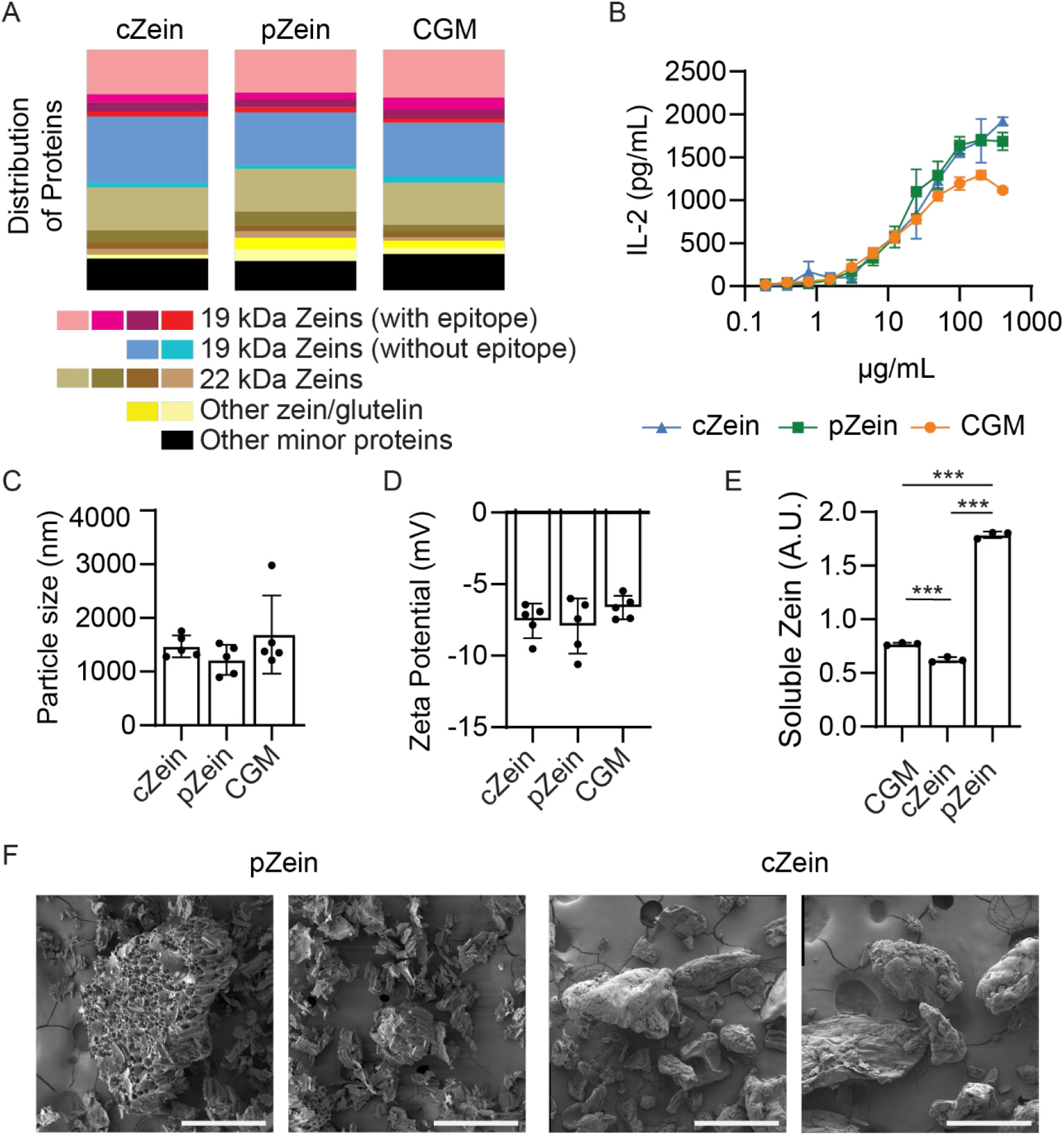
Biochemical and biophysical data on zein preparations. **(A)** Distribution of zein isoforms and other proteins identified in proteomics analysis between cZein, pZein, and CGM. **(B)** In a co-culture system with primary APCs and zein-specific T cell heterologous lines, cZein, pZein, and CGM are all equally stimulatory (N=3/group). **(C)** Particle size between cZein, pZein, and CGM in SIF (N=5/group). **(D)** Zeta potential between cZein, pZein, and CGM in SIF (N=5/group). **(E)** Solubility of CGM, cZein, and pZein in SIF (N=3/group). **(F)** SEM images of cZein and pZein. Scale bars = 500 µm. * P < 0.05, ** P < 0.01, *** P < 0.001. Error bars represent standard deviation.

### Development of zein-specific T cells is limited by intestinal uptake

Armed with structural and solubility differences, we sought to determine how each zein preparation interfaces with the intestinal immune system, to better understand differences in capacity to induce a zein-specific T cell response. We reasoned that induction of an intestinal T cell population may broadly rely on three steps: (1) antigen uptake across the intestinal epithelium and sampling by antigen-presenting cells (APCs); (2) activation of the APCs and trafficking to the draining lymph node; and (3) activation and differentiation of naïve T cells and their return to establish intestinal residency. Based on data from food allergy models, we first tested the APC activation hypothesis. As shown above, different zein preparations were equally stimulatory in an *in vitro* T cell activation assay, providing some evidence that antigen presentation does not differentiate the zein preparations (**Fig 2B**). As further confirmation, we stimulated isolated CD11c+ APCs with each zein preparation. Two markers of APC activation, CD80 and CD86, were only weakly induced compared to an LPS control, and not in a pattern mirroring intestinal T cell induction (**Fig S3A-D**). Further suggesting that APC activation is not the limiting step for zein-specific T cell induction, APCs isolated from mice consuming AAD diet showed higher activation marker levels compared to mice on chow diets (**Fig S3E-H**). These data highlight that a more biochemically complex diet (chow) does not globally support APC activation. Taken together, these data suggest that APC activation is unlikely to explain differences in development of a T cell population following feeding of cZein, pZein, or CGM.

Since differential APC activation was not observed, we next considered that zein preparations may differ in sampling across the intestinal epithelium, thereby controlling antigen availability. To determine whether differences in intestinal uptake contribute to zein-specific T cell induction, we needed an assay sensitive enough to detect unmodified protein, since fluorophore labeling could alter the biochemical properties of zein that control uptake. Initial attempts to quantify zein in serum or intestinal tissue did not provide sufficient sensitivity or suitable controls to reliably quantify intestinal uptake. We reasoned that intestinal uptake or digestion would reduce the amount of antigen recovered in feces. Our assay “fecal recovery after gavage (FRAG)” measures antigen transit into feces following gavage with a standardized protein dose (**Fig 3A**). To validate assay sensitivity, we confirmed that a drug previously shown to inhibit intestinal OVA uptake (zileuton)^23^, increased fecal OVA recovery in the FRAG assay (**Fig S4A**). For zein preparations, FRAG revealed high fecal recovery for cZein, suggesting limited uptake, while pZein and CGM were depleted in feces, consistent with either intestinal uptake or digestion (**Fig 3B**). As additional evidence that intestinal uptake contributes to the differential response to zein preparations, when cZein, pZein, or CGM were delivered as a subcutaneous injection that bypasses the intestinal epithelium, T cell induction was comparable between the preparations (**Fig S4B**). Together, these data suggested that limited intestinal uptake could explain the reduced intestinal T cell response toward cZein compared to pZein and CGM.

**Figure 3:**
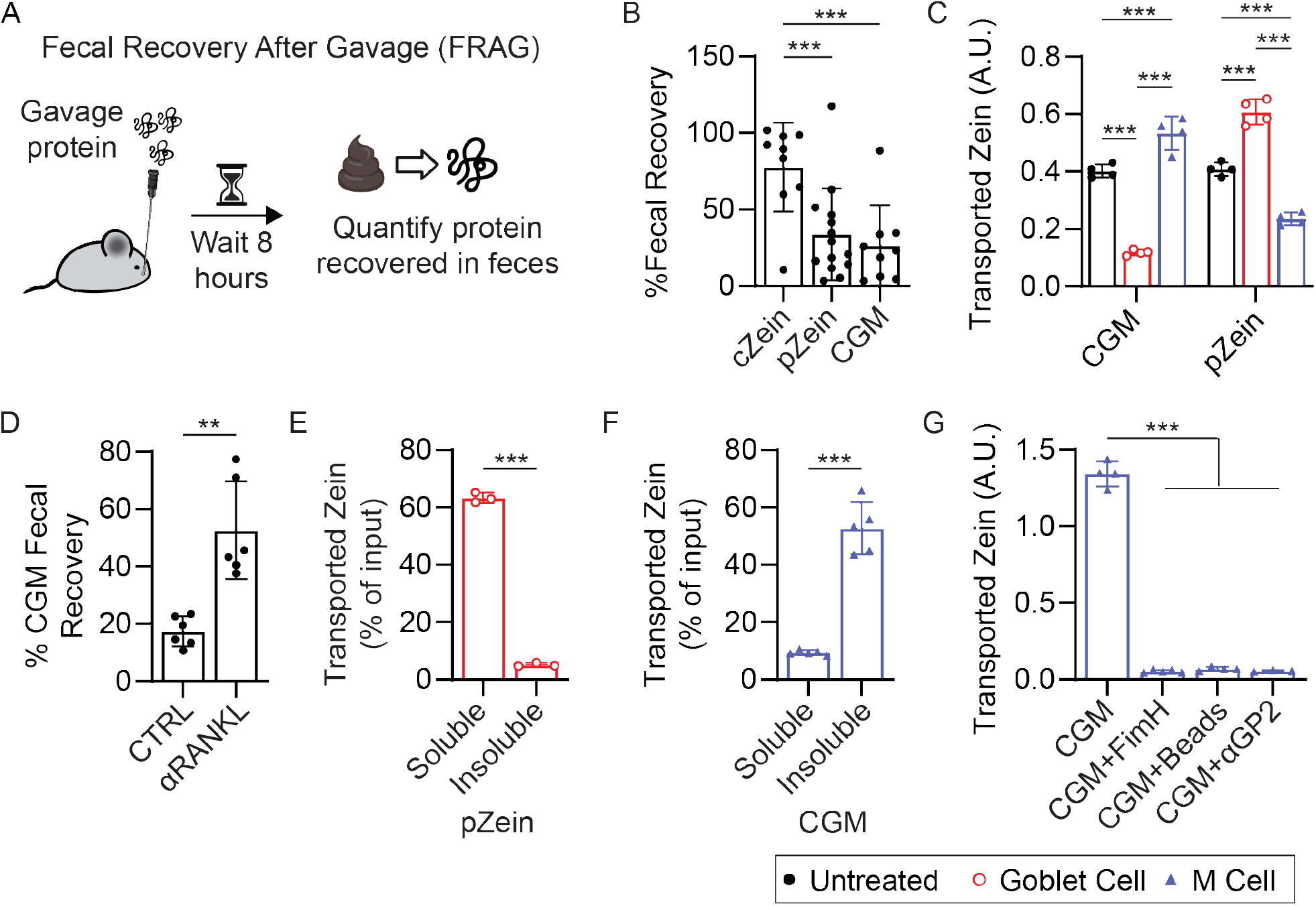
Intestinal uptake differentiates zein preparations. **(A**) The fecal recovery after gavage (FRAG) assay uses fecal antigen recovery as a readout for host accessibility. **(B)** Fecal recovery of zein following gavage with cZein, pZein, or CGM (N=9-14/group). **(C)** Transport of CGM and pZein through goblet and M cells (N=4/group). **(D)** Fecal recovery of zein in mice injected with αRANKL or CTRL antibody and then gavaged with CGM. **(E)** Transport of the soluble and insoluble pZein fraction through goblet cells (N=3/group). **(F)** Transport of the soluble and insoluble CGM fraction through M cells (N=5/group). **(G)** Transport of CGM through M cells when added alone or alongside FimH, beads, or αGP2 antibody (N=3-5/group). * P < 0.05, ** P < 0.01, *** P < 0.001. Error bars represent standard deviation.

Next, we evaluated mechanisms of intestinal zein uptake. As a prolamin protein, zein has limited solubility in aqueous solutions, which may influence intestinal uptake. As previously noted in our SIF solubility assay, pZein was over twice as soluble as cZein (**Fig 2E**). Since pZein and CGM had different apparent zein solubilities yet showed similar recovery in our FRAG assay, solubility alone did not appear to explain their intestinal handling. We therefore considered whether the zein preparations could differ in the route through which they were sampled. Two major antigen sampling mechanisms have been described in the intestine: goblet cells and M cells. Goblet cell antigen sampling through goblet-cell associated antigen passages (GAPs) is the best described food antigen sampling route, based on data using OVA^24,25^. M cells are better known for their role in sampling pathogenic bacteria, but also hypothesized to transport food antigens^26^. To evaluate uptake route, we developed a polarized intestinal organoid assay that combines established transwell culture methods with differentiation protocols for goblet and M cells^27,28^. Organoids were seeded onto transwell inserts, where they form an apical luminal surface and a basolateral membrane. Adding DAPT stimulated the differentiation of goblet cells, while a combination of RANKL and TNFα induced M cell development. We validated the selective induction of goblet and M cells using established ligands (**Fig S4C-E**). We found that CGM is transported by M cells, while pZein is transported by goblet cells, confirming different uptake routes for different zein preparations (**Fig 3C**). Our observation that pZein is a goblet cell ligand is consistent with other soluble food antigens. However, few dietary M cell ligands have been described. To validate that CGM is an M cell ligand *in vivo*, we depleted M cells by injecting mice with an αRANKL antibody^29^. FRAG analysis showed that mice treated with αRANKL had significantly more zein recovered in feces following CGM gavage compared to mice injected with the control antibody (**Fig 3D**). Thus, consistent with our organoid findings, these data support a role for M cells in CGM-associated zein sampling *in vivo*. Since neither pZein nor CGM are fully soluble in organoid media, we next evaluated whether uptake route tracked with solubility. As expected, pZein was primarily transported through goblet cells in the soluble state (**Fig 3E**). Conversely, CGM was primarily sampled as an insoluble antigen (**Fig 3F**). M cells transport bacteria through the GP2 receptor binding the bacterial adhesion molecule FimH, and particulates are transported through receptor-independent endocytosis^30^. Thus, to gain insight into the mechanisms of M cell-mediated zein transport, we co-exposed M cells to CGM alongside soluble FimH, latex beads, or an αGP2 antibody. M cell-mediated transport of CGM was inhibited in all cases (**Fig 3G**). Thus, our data suggest that CGM may be transported through mechanisms involving GP2, and converging on a transcytosis pathway shared with other particulate ligands.

Taken together, these data show that zein can be sampled by goblet or M cells depending on the source of the protein. Because CGM feeding induces zein-specific Tregs, these findings demonstrate that M cell-mediated antigen transport is compatible with the development of an intestinal Treg response. Further, manipulating M cell abundance is sufficient to change zein uptake, as measured by residual protein in fecal samples and an *in vitro* transwell assay.

### Intestinal uptake can be modified by food formulation

We suspected that solubility explains the increased intestinal uptake and more robust zein-specific T cell induction in response to pZein compared to cZein. To explicitly test this, we used a saccharide conjugation protocol previously shown to increase zein solubility in aqueous systems^31^. As expected, soluble zein (SOL-cZein) was significantly more soluble than control zein (CTRL-cZein), but did not differ in particle size and charge, or in T cell activation in our *in vitro* assay (**Fig 4A-C, Fig S5A-B**). FRAG analysis showed less fecal zein recovery for SOL-cZein, consistent with increased intestinal uptake (**Fig 4D**). Furthermore, our organoid model revealed that SOL-cZein was transported by goblet cells, consistent with pZein, and as expected for a soluble dietary antigen (**Fig 4E**). Consistent with the decreased fecal recovery and increased goblet cell transport, feeding a diet containing SOL-cZein induced significantly more zein-specific intestinal T cells compared to CTRL-cZein (**Fig 4F**). These data show that manipulating the physical properties of an antigen is sufficient to modify intestinal uptake, controlling the development of an ensuing cognate T cell response.

**Figure 4:**
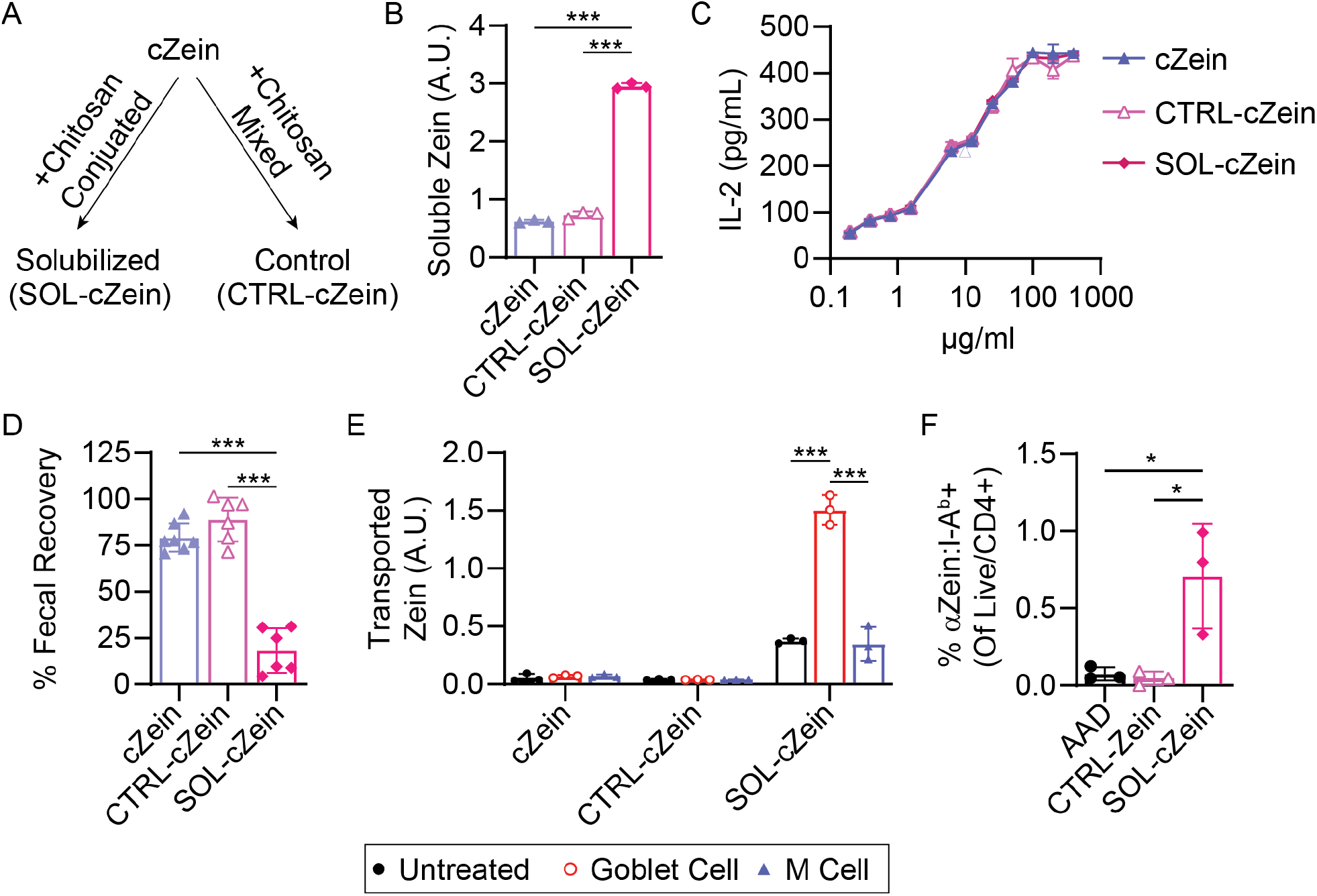
Saccharide conjugation to cZein improves solubility and drives goblet cell mediated uptake. **(A)** cZein was solubility-enhanced (SOL-cZein) or incubated in a mock reaction (CTRL-cZein). **(B)** SOL-cZein showed increased solubility in SIF compared to cZein or CTRL-cZein (N=3/group). **(C)** In a co-culture system with primary APCs and zein-specific T cell heterologous lines, cZein, CTRL-cZein, and SOL-cZein are all equally stimulatory (N=3/group). **(D)** Fecal recovery of zein following gavage with cZein, CTRL-cZein, or SOL-cZein (N=6-7/group). **(E)** Transport of cZein, CTRL-cZein, and SOL-cZein through goblet and M cells (N=3/group). **(F)** Intestinal zein-specific T cells following feeding with AAD, CTRL-cZein and SOL-cZein. * P < 0.05, ** P < 0.01, *** P < 0.001. Error bars represent standard deviation.

Our data suggest that intestinal uptake can be a limiting determinant of T cell induction and is tunable by food formulation. To determine the generalizability of this finding, we focused on peanut. Roasting peanut is known to change biophysical and biochemical properties, and can increase allergenicity, providing another example of a difference in protein or food format leading to a difference in immune cell recognition^32^. Consistent with prior reports, roasted peanut showed the anticipated decrease in solubility of the allergen Ara h 1 compared to raw peanut in organoid media, the conditions used in our transport assay; however, in our study, roasted peanut unexpectedly showed slightly increased solubility in SIF (**Fig 5A**). Roasting peanut did not change measured particle size or zeta potential (**Fig S5C-D**). Using our organoid assay, we found that Ara h 1 from raw peanuts is predominantly transported by goblet cells, while Ara h 1 from roasted peanuts showed more M cell translocation *in vitro* (**Fig 5B**). Further, inhibiting M cell development *in vivo* increased fecal recovery of Ara h 1 following gavage with roasted peanut, confirming that when delivered in roasted peanut, Ara h 1 is an M cell ligand (**Fig 5C**). Thus, beyond zein, the same protein can be sampled by different routes depending on protein form.

**Figure 5:**
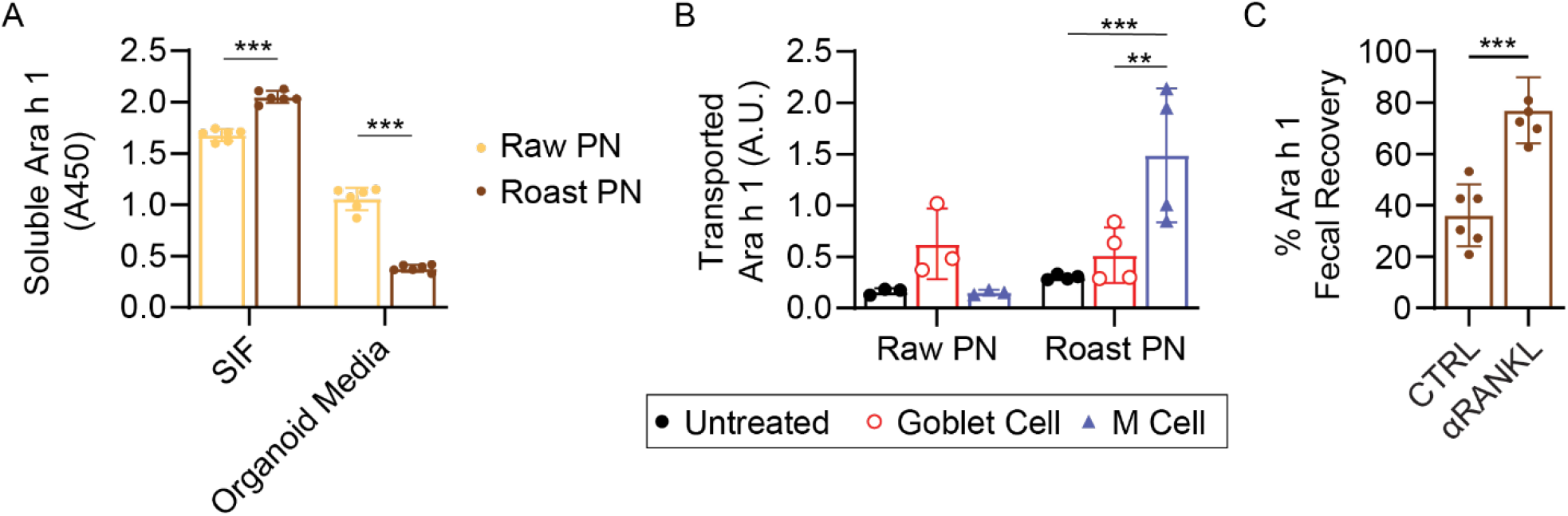
Roasting peanut turns Ara h 1 into an M cell ligand. **(A)** Solubility of Ara h 1 in SIF and organoid media measured by ELISA (N=6/group). **(B)** Transport of Ara h 1 through goblet cells and M cells (N=3-4/group). **(C)** Fecal recovery of Ara h 1 in mice injected with αRANKL or CTRL antibody and then gavaged with roasted peanut.* P < 0.05, ** P < 0.01, *** P < 0.001. Error bars represent standard deviation.

## Discussion

Our data suggest that epithelial uptake is a limiting determinant of intestinal CD4 T cell responses. Antigen-specific T cell development varied 10-fold in response to different preparations of the same protein. Context-dependent T cell responses are usually explained by adjuvant molecules that activate APCs, and factors that influence the extent and route of antigen sampling are far less resolved. Goblet cell antigen sampling through goblet cell associated passages (GAPs) has emerged as the dominant pathway for food antigen sampling, based largely on observations from soluble antigens^33^. Similarly, solubilized zein is transported by goblet cells, while certain insoluble zein preparations are preferentially sampled by M cells. Increasing zein solubility was sufficient to promote sampling by goblet cells, increase intestinal uptake, and stimulate zein-specific T cell development. While GAPs are established to vary depending on microbial signals and EGF levels in breast milk^25,33^, intestinal or food-derived factors that influence M cell sampling are less established.

The role of antigen uptake in determining the magnitude of the T cell response toward zein aligns with multiple recent studies showing that intestinal sampling controls immune responses toward food proteins. For example, recent work suggests that cysteinyl leukotriene-mediated goblet cell antigen sampling is the central determinant of susceptibility to oral anaphylaxis across different mouse strains^23^. Further, food allergy is associated with antigen sampling through secretory epithelial cells^34^. Notably, while these examples focus on intestinal antigen uptake as a determinant of food allergy, our data suggest that differences in intestinal antigen uptake can also shape the magnitude of antigen-specific responses associated with oral tolerance.

While M cells have been postulated to play a role in food antigen sampling, few dietary M cell ligands have been described^26,35^. In one example, the milk protein casein was observed to accumulate in Peyer’s patches following gavage, suggesting an M cell-mediated transport pathway^36^. Notably, the soluble milk antigens alpha- and beta-lactoglobulin only accumulated in Peyer’s patches following pasteurization, a treatment that reduced their solubility. Two potential mechanisms could explain M cell-mediated antigen sampling. First, particle shape or other physical properties may promote M cell sampling. Studies with nanoparticles have revealed surface hydrophobicity, rigidity, and shape as determinants of epithelial uptake^37,38^. Alternatively, our FimH and αGP2 co-incubation experiments suggest that CGM may be transported through M cells via GP2. It is possible that an insoluble fiber or other CGM component is the direct GP2 ligand, and that zein can be co-transported when part of the same food particle, but not in response to simple co-exposure. Plant non-starch polysaccharides were previously shown to block M cell transport of *E. Coli*, though whether the effect was mediated by competition for uptake was not explored^39^. Either a biophysical difference or direct GP2 ligand could explain why CGM, but not cZein, which was similarly insoluble, can be transported by M cells.

The dependence of food-specific T cell development on the microbiome varies between antigens. For example, germ-free mice have fewer zein-specific intestinal T cells compared to colonized mice^16^. A recent study reported food antigen-specific populations that differed in their dependence on the microbiome, though the dietary antigens were not mapped, limiting insights into antigen features that may predict microbe dependency^40^. While the role of the microbiome in global immune system maturation is a possible mediator, it is unclear why this effect would only manifest for some food antigens^41^. Alternatively, the microbiome may influence T cell responses to food antigens by modulating intestinal uptake. While pathogenic microbes are known to stimulate M cells, emerging evidence suggests that certain microbial communities or pasteurized commensals can also increase M cell maturation^42–45^. If gut microbes selectively modulate antigen sampling pathways, M cell dependence could explain why some dietary antigens appear microbiome dependent, while others are independent.

Within the past 2 years, RORγt-expressing APCs have been shown to be critical for intestinal Treg induction toward microbial and food proteins, including zein^16,46–48^. However, the coordination of intestinal antigen uptake and antigen presentation is not well explored. It is unclear whether goblet and M cells directly pass dietary antigens to RORγt APCs through a coordinated hand-off, whether antigens are “dumped” at the basolateral membrane and then separately sampled by RORγt APCs, or whether a third cell type may mediate the exchange. While lymph node trafficking is essential for immune tolerance to soluble antigens sampled through goblet cells, the site of T cell presentation for M cell-sampled antigens remains unresolved^49^. Notably, human M cells were recently shown to express MHCII and directly present gliadin antigens to CD4 T cells; however, MHCII expression was not detected in mouse M cells in the same study, suggesting that direct antigen presentation may be a human-specific M cell function^50^. Thus, in our mouse model, M cell-sampled zein likely requires transfer to a professional APC before T cell activation.

Overall, we revealed an antigen for which intestinal sampling is the apparent limiting step for intestinal T cell induction, and showed that different forms of the same protein can be sampled through distinct intestinal routes. Together, these findings identify antigen uptake as a regulated step that can shape the magnitude of the Treg response following dietary exposure. This work further establishes physiologically relevant M cell ligands associated with immune tolerance. Further, these data suggest that the same antigen can be sampled by both goblet and M cells, with the dominant route likely dependent on physical or biochemical properties of the antigen or food matrix. Our findings underscore that antigen sampling is a regulated step of T cell programming and suggest that altering the route of uptake could influence immune interpretation of food antigens.

## Acknowledgements

The authors thank the NIH tetramer core facility for providing MHCII tetramer reagents. JEB was an HHMI awardee of LSRF. This work was supported by the Flow Cytometry Core Facility of the Salk Institute (RRID:SCR_014839) with funding from NIH-NCI CCSG: P30 CA014195. Data was collected on instruments in the Shared FACS Facility obtained using NIH S10 Shared Instrument Grants (S10RR027431-01 and 1S10OD023831-01). This work was supported by the Waitt Advanced Biophotonics Core Facility of the Salk Institute (RRID:SCR_014838) with funding from NIH-NCI CCSG P30 CA014195, NIH-NIA San Diego Nathan Shock Center P30 AG068635, and the Waitt Foundation. We thank Chynna Bowman and Daniela Boassa for their assistance with SEM imaging. This work was supported by the Mass Spectrometry Core of the Salk Institute (RRID:SCR_014843) with funding from NIH-NCI CCSG P30 CA014195, NIH-NIA San Diego Nathan Shock Center P30 AG068635, two NIH Shared Instrumentation Grants S10-OD021815 (ThermoFisher Q-Exactive quadrupole orbitrap) and S10-OD038262 (ThermoFisher Orbitrap IQ-X tribrid), and the Helmsley Center for Genomic Medicine. Thanks to Elizabeth Sattely for helpful discussions and manuscript feedback.

## Author Contributions

**Conceptualization:** JEB, YC, HHL; **Investigation:** YC, HHL, AM, RK, ES, TC, JEB; **Formal Analysis:** YC, HHL, JEB; **Writing-original draft:** JEB; **Writing-review & editing:** YC, HHL, RK, JEB; **Supervision:** JEB. All authors reviewed and approved the final manuscript.

## References

1. Tordesillas, L. & Berin, M. C. Mechanisms of Oral Tolerance. Clin. Rev. Allergy Immunol. 55, 107–117 (2018).

2. Pabst, O. & Mowat, A. M. Oral tolerance to food protein. Mucosal Immunol. 5, 232–239 (2012).

3. Hadis, U. et al. Intestinal Tolerance Requires Gut Homing and Expansion of FoxP3+ Regulatory T Cells in the Lamina Propria. Immunity 34, 237–246 (2011).

4. Chen, Q., Abdi, A. M., Luo, W., Yuan, X. & Dent, A. L. T follicular regulatory cells in food allergy promote IgE via IL-4. JCI Insight 9, e171241 (2024).

5. Gowthaman, U. et al. Identification of a T follicular helper cell subset that drives anaphylactic IgE. Science 365, eaaw6433 (2019).

6. Steele, L., Mayer, L. & Cecilia Berin, M. Mucosal immunology of tolerance and allergy in the gastrointestinal tract. Immunol. Res. 54, 10.1007/s12026-012-8308-4 (2012).

7. Tordesillas, L. et al. Skin exposure promotes a Th2-dependent sensitization to peanut allergens. Journal of Clinical Investigation 124, 4965–4975 (2014).

8. Abdel-Gadir, A. et al. Microbiota therapy acts via a regulatory T cell MyD88/RORγt pathway to suppress food allergy. Nat. Med. 25, 1164–1174 (2019).

9. Tamura, S. ichi, Shoji, Y., Hasiguchi, K., Aizawa, C. & Kurata, T. Effects of cholera toxin adjuvant on IgE antibody response to orally or nasally administered ovalbumin. Vaccine 12, 1238–1240 (1994).

10. Dunkin, D., Berin, M. C. & Mayer, L. Allergic sensitization can be induced via multiple physiologic routes in an adjuvant-dependent manner. Journal of Allergy and Clinical Immunology 128, 1251–1258.e2 (2011).

11. Zhang, T. et al. Boiling and roasting treatment affecting the peanut allergenicity. Ann. Transl. Med. 6, 357 (2018).

12. Abbring, S. et al. Milk processing increases the allergenicity of cow’s milk—Preclinical evidence supported by a human proof-of-concept provocation pilot. Clinical and Experimental Allergy 49, 1013 (2019).

13. Tan, J. et al. Dietary Fiber and Bacterial SCFA Enhance Oral Tolerance and Protect against Food Allergy through Diverse Cellular Pathways. Cell Rep. 15, 2809–2824 (2016).

14. Li, W. et al. A bacterial bile acid metabolite modulates Treg activity through the nuclear hormone receptor NR4A1. Cell Host Microbe 29, 1366–1377.e9 (2021).

15. Nagai, M. et al. Sugar and arginine facilitate oral tolerance by ensuring the functionality of tolerogenic immune cell subsets in the intestine. Cell Rep. 43, 114490 (2024).

16. Blum, J. E. et al. Identification and characterization of dietary antigens in oral tolerance. Sci. Immunol. 11, (2026).

17. Kim, K. S. et al. Dietary antigens limit mucosal immunity by inducing regulatory T cells in the small intestine. Science 351, 858–863 (2016).

18. Nagai, M. et al. Sugar and arginine facilitate oral tolerance by ensuring the functionality of tolerogenic immune cell subsets in the intestine. Cell Rep. 43, (2024).

19. Anderson, T. J. & Lamsa, B. P. Zein extraction from corn, corn products, and coproducts and modifications for various applications: A review. Cereal Chem. 88, 159–173 (2011).

20. Foged, C., Brodin, B., Frokjaer, S. & Sundblad, A. Particle size and surface charge affect particle uptake by human dendritic cells in an in vitro model. Int. J. Pharm. 298, 315–322 (2005).

21. Oyewumi, M. O., Kumar, A. & Cui, Z. Nano-microparticles as immune adjuvants: correlating particle sizes and the resultant immune responses. Expert Rev. Vaccines 9, 1095–1107 (2010).

22. Giteru, S. G., Ali, M. A. & Oey, I. Recent progress in understanding fundamental interactions and applications of zein. Food Hydrocoll. 120, 106948 (2021).

23. Hoyt, L. R. et al. Cysteinyl leukotrienes stimulate gut absorption of food allergens to promote anaphylaxis in mice. Science 389, (2025).

24. Gustafsson, J. K. et al. Intestinal goblet cells sample and deliver lumenal antigens by regulated endocytic uptake and transcytosis. Elife 10, (2021).

25. Knoop, K. A., McDonald, K. G., McCrate, S., McDole, J. R. & Newberry, R. D. Microbial Sensing by Goblet Cells Controls Immune Surveillance of Luminal Antigens in the Colon. Mucosal Immunol. 8, 198 (2014).

26. Kulkarni, D. H. & Newberry, R. D. Antigen Uptake in the Gut: An Underappreciated Piece to the Puzzle? 32, 5 (2026).

27. Yin, X. et al. Niche-independent high-purity cultures of Lgr5+ intestinal stem cells and their progeny. Nat. Methods 11, 106–112 (2014).

28. Kanaya, T. et al. Development of intestinal M cells and follicle-associated epithelium is regulated by TRAF6-mediated NF-κB signaling. J. Exp. Med. 215, 501–519 (2018).

29. Knoop, K. A. et al. RANKL is necessary and sufficient to initiate development of antigen-sampling M cells in the intestinal epithelium. J. Immunol. 183, 5738–5747 (2009).

30. Ohno, H. & Hase, K. Glycoprotein 2 (GP2): Grabbing the FimH+ bacteria into M cells for mucosal immunity. Gut Microbes 1, 407 (2010).

31. Wang, X. J. et al. Preparation of glycosylated zein and retarding effect on lipid oxidation of ground pork. Food Chem. 227, 335–341 (2017).

32. Maleki, S. J., Chung, S. Y., Champagne, E. T. & Raufman, J. P. The effects of roasting on the allergenic properties of peanut proteins. Journal of Allergy and Clinical Immunology 106, 763–768 (2000).

33. Kulkarni, D. H. et al. Goblet Cell Associated Antigen Passages Support the Induction and Maintenance of Oral Tolerance. Mucosal Immunol. 13, 271 (2019).

34. Noah, T. K. et al. IL-13–induced intestinal secretory epithelial cell antigen passages are required for IgE-mediated food-induced anaphylaxis. Journal of Allergy and Clinical Immunology 144, 1058–1073.e3 (2019).

35. Reichel, P. E., Chakraborty, S., Tay, H. L. & Hogan, S. P. Dietary Antigen Interaction with Intestinal Epithelial Cells. Current Allergy and Asthma Reports 2026 26:1 26, 13-(2026).

36. Roth-Walter, F. et al. Pasteurization of milk proteins promotes allergic sensitization by enhancing uptake through Peyer’s patches. Allergy 63, 882–890 (2008).

37. Pridgen, E. M., Alexis, F. & Farokhzad, O. C. Polymeric Nanoparticle Drug Delivery Technologies for Oral Delivery Applications. Expert Opin. Drug Deliv. 12, 1459 (2015).

38. Zheng, Y. et al. Transepithelial transport of nanoparticles in oral drug delivery: From the perspective of surface and holistic property modulation. Acta Pharm. Sin. B 14, 3876 (2024).

39. Roberts, C. L. et al. Translocation of Crohn’s disease Escherichia coli across M-cells: contrasting effects of soluble plant fibres and emulsifiers. Gut 59, 1331–1339 (2010).

40. Yi, J. et al. A hierarchy of intestinal antigens instructs the CD4+ T cell receptor repertoire. Immunity 58, 1217–1235.e4 (2025).

41. Belkaid, Y. & Hand, T. W. Role of the Microbiota in Immunity and inflammation. Cell 157, 121 (2014).

42. Zhang, W. et al. Pasteurized Akkermansia muciniphila promotes GP2 expression in microfold cells and facilitates Salmonella infection. Protein Cell 10.1093/PROCEL/PWAF017 (2025) doi:10.1093/PROCEL/PWAF017.

43. Donaldson, D. S., Pollock, J., Vohra, P., Stevens, M. P. & Mabbott, N. A. Microbial Stimulation Reverses the Age-Related Decline in M Cells in Aged Mice. iScience 23, (2020).

44. Alfituri, O. A. et al. Differential role of M cells in enteroid infection by Mycobacterium avium subsp. paratuberculosis and Salmonella enterica serovar Typhimurium. Front. Cell. Infect. Microbiol. 14, 1416537 (2024).

45. Tahoun, A. et al. Salmonella Transforms Follicle-Associated Epithelial Cells into M Cells to Promote Intestinal Invasion. Cell Host Microbe 12, 645–656 (2012).

46. Fu, L. et al. PRDM16-dependent antigen-presenting cells induce tolerance to gut antigens. Nature 2025 642:8068 642, 756–765 (2025).

47. Cabric, V. et al. A wave of Thetis cells imparts tolerance to food antigens early in life. Science (1979). 389, 268–274 (2025).

48. Rodrigues, P. F. et al. Rorγt-positive dendritic cells are required for the induction of peripheral regulatory T cells in response to oral antigens. Cell 188, 2720–2737.e22 (2025).

49. Worbs, T. et al. Oral tolerance originates in the intestinal immune system and relies on antigen carriage by dendritic cells. J. Exp. Med. 203, 519 (2006).

50. Wang, D. et al. Human gut M cells resemble dendritic cells and present gluten antigen. Nature 2025 650:8100 650, 251–260 (2025).

