## Supplemental Information for "Intestinal uptake regulates T cell responses to dietary antigens"

### Materials and Methods

#### Animal Experiments

All procedures with animals were approved by the Salk Institute or Stanford University institutional animal care and use committees and adhered to ethical standards. For zein-specific T cell induction studies, C57BL/6 mice were bred onto L-AA Defined AIN93G (AAD, Dyets) diet and exposed to diets containing fractions of interest for two weeks, starting between 3 and 8 weeks of age. Studies where mice were bred onto diets use male and female mice equally. For studies involving chow mice, we purchased 6-week-old C57Bl/6 mice from Jackson Laboratory (Strain: 000664). In the OTII adoptive transfer experiment, $Ptprc^{a}Pepc^{b}$/BoyJ mice (Jackson Laboratory) were retrooribtally injected with 1E6 OTII cells from the spleen of a donor mouse (Strain 11490, Taconic).

cZein was zein from Santa Cruz biotechnology, specifically the 500 g package size (sc-216069). The 1kg package size had different biochemical properties and was not included in this study. pZein was from FloZein Enterprises. CGM was obtained from both FloZein Enterprises and Teklad. spentCGM was from FloZein Enterprises. OVA and gliadin were obtained from Sigma. Custom diets were prepared in lab by mixing L-AA Defined AIN93G diet powder (Dyets) with proteins or food fractions of interest to the indicated percentage (1% or 10%). Mice had *ad libitum* access to food and water for the duration of the study. In experiments with peanut, raw peanuts were roasted in lab at 320°F for 50 minutes.

In one experiment, mice were injected with zein fractions emulsified in CFA (InvivoGen). In another experiment, mice were injected every other day with αRANKL (bioXCell, BE0191) or control (bioXCell, BE0089) antibodies as previously described[1]. In one experiment mice were gavaged with 100 mg/kg zileuton (Cayman Chemical) in 0.5% methylcellulose, as previously described[2].

#### Intestinal T cell Isolation

The small intestine was removed from mice and cleared of adipose tissue and Peyer's patches. The intestine was then cut into segments, a sagittal cut was used to open the intestine, and the sample was incubated in DMEM (Corning) + 10% fetal bovine serum (FBS, Avantor) with 5 mM EDTA (Chem Impex) and 1 mM DTT (Sigma) at 37°C with gentle shaking for 40 minutes. The solution was passed over a filter to remove liquid, the intestine was placed into a 50 ml conical tube with 20 ml of DMEM, and the tube was vigorously shaken for 15 seconds. This washing procedure was then repeated. The intestine was then transferred to a GentleMACS C tube (Miltenyi Biotec) and digested on the OctoMACS Tissue Dissociator using the mouse lamina propria dissociation kit (Miltenyi Biotec). After dissociation, the solution was mixed 1:1 with 80% Percoll (Cytiva), moved to a 15 ml tube, and layered over an 80% percoll layer. The tubes were centrifuged at 600 x g for 10 minutes. The interface layer was then transferred to a fresh tube and washed with DMEM + 10% FBS to remove residual percoll.

#### Tetramer Staining and Flow Cytometry

MHCII tetramers targeting zein, OVA, and gliadin were obtained from the NIH Tetramer Core Facility. They were used at either a dilution of 1:100 in RPMI (Corning) + 10% FBS for 1 h at 37°C or room temperature at 1:200 in MACS Buffer (Miltenyi Biotec) + 2% BSA for 1 hour. These staining protocols gave equivalent signal.

Cells were stained for 30 minutes at room temperature with viability dye (GhostDye Red 780, Cytek) and in some experiments cell surface markers. If transcription factor staining was needed, cells were fixed (eBiosciences Foxp3/Transcription Factor Kit) overnight at 4°C, permeabilized, and stained the following morning. Data were collected on an LSR (BD Bioscience) flow cytometer.

The antibodies used in this study include: Brilliant Violet 711 anti-mouse CD4 (Clone RM4-5, Biolegend 100549), Brilliant Violet 510 anti-mouse CD4 (Clone RM4-5, Biolegend, 100553), PE/Cyanine7 anti-mouse Foxp3 (Clone FJK-16S, Thermo Fisher, 25-5773-82), APC anti-mouse CD45.2 (Clone 104, Biolegend, 10981), PE anti-mouse CD45.1 (Clone A20, Biolegend, 110707), PE-Dazzle anti-mouse TCRβ (Clone H57-597, Biolegend, 109239), AF700 anti-mouse TCRβ (Clone H57-597, BD, 560705), BV421 anti-mouse I-A/I-E (Clone M5/114.15.2, Biolegend, 107631), BV605 anti-mouse CD11c (Clone N418, Biolegend, 117333), PerCP-Cy5.5 anti-mouse CD80 (Clone 16-10A1, Biolegend, 104721), FITC anti-mouse CD86 (Clone A17199A, Biolegend, 159219), PE/Cy5 anti-mouse CD86 (Clone GL-1, Biolegend, 105015), BV605 anti-mouse CD80 (Clone 16-10A1, Biolegend, 104729).

**Co-culture systems with APCs and T cell Transgenic Lines**
The T cell transgenic lines specific for zein were developed in our prior work[3]. To indue dendritic cell expansion, mice were injected with 5 x $10^6$ B16-FLT3L cells. Spleens were harvested and dendritic cells were isolated using the spleen dissociation kit and CD11c microbeads UltraPure kit following manufacturer’s instructions (Miltenyi Biotec, #130-095-926 and #130-125-835). Co-cultures used 50,000 dendritic cells and 20,000 TCR transgenic cell lines in media containing DMEM (Corning) + 10% FBS (Avantor) + 1% penicillin-streptomycin (Gibco) + 1% Glutamax (Gibco). After 24 hours of culture, media IL2 levels were measured using ELISA assays (Biolegend). To measure CD80 and CD86, freshly isolated CD11c+ dendritic cells were incubated alone (no T cells) for 18 hours with 50 ug/ml of zein fractions or 5 ug/ml LPS (eBioscience).

**Fecal Recovery After Gavage Assay**
Mice were orally gavaged with 5 mg of food antigen suspended in 200 µL of 5% bicarbonate buffer. Eight hours after gavage, fecal pellets were collected and lyophilized. As an input control, a weight matched fecal pellet sample from mice on antigen-free diet was prepared and mixed with 5 mg food antigen. Lyophilized fecal samples were resuspended in an extraction volume of 1 mL 1% SDS per 100 mg of feces. Samples were homogenized by bead beating and centrifuged. The resulting supernatants were collected for antigen quantification. For fecal antigen quantification, high-binding 96-well plates (Corning) were coated with 100 µL of fecal extract per well and incubated overnight at 4 °C. Fecal antigens were detected using antigen-specific antibodies against 19-kDa α-zein (MyBioSource), ovalbumin (Novus Biologicals), or Ara h 1 (MyBioSource). Each sample was normalized to its weight matched input control.

**Solubility Assays**
Food antigens were prepared at a concentration of 1 mg/mL in either Simulated Intestinal Fluid (SIF) buffer (RICCA Chemical Company) or IntestiCult™ Organoid Growth Medium (STEMCELL Technologies), incubated on an end-over-end rotator at room temperature for 4 h, and centrifuged to recover the soluble fraction. For quantification, high-binding 96-well plates were coated with 100 µL of supernatant per well and incubated overnight at 4 °C. Antigens were detected using antigen-specific antibodies against 19-kDa α-zein (MyBioSource) or Ara h 1 (MyBioSource).

**Organoid Maintenance**
Mouse intestinal organoids were cultured as domes containing 50% Matrigel (STEMCELL Technologies) and 50% Mouse Organoid Growth Medium (STEMCELL Technologies). Organoid domes were maintained in mouse organoid medium supplemented with 100 µg/mL penicillin/streptomycin in 24-well plates at 37 °C with 5% $CO_2$. Culture medium was refreshed

every 2–3 days. Organoids were passaged when the culture density reached approximately 150 organoids per well.

**Organoid Transwell Assays**

Organoids were harvested using Gentle Cell Dissociation Reagent (STEMCELL Technologies) and incubated at room temperature for 15 min with shaking. An equal volume of DMEM/F-12 (STEMCELL Technologies) was then added, and the organoids were washed by centrifugation at 200 × *g* for 5 min at 4 °C. To generate organoid-derived monolayers, 6.5-mm Transwell inserts with 0.4-µm pores were pre-coated with 2% Matrigel in D-PBS. The harvested organoids (~300) were seeded into the apical compartment in Human Organoid Differentiation Medium (STEMCELL Technologies) supplemented with 10 µM Y-27632 (STEMCELL Technologies) and 100 µg/mL penicillin/streptomycin. The apical and basolateral compartments contained 100 µL and 500 µL of medium, respectively. For M-cell differentiation, organoid-derived monolayers were treated with 500 ng/mL RANKL (Gibco) and 10 ng/mL TNF-α (Gibco) for 72 h. For goblet-cell differentiation, monolayers were treated with 10 µM DAPT (Thermo Scientific Chemicals) for 4 days. During goblet-cell differentiation, culture medium was replaced every 48 h. For antigen transport assays, medium from both the apical and basolateral compartments was removed and replaced with fresh organoid medium. Food antigens were added to the apical compartment at a final concentration of 500 µg/mL. For microsphere transport assays, FluoSpheres Microspheres (Invitrogen) were added at a 1:250 dilution. After incubation for 4 h at 37 °C, basolateral medium was collected for antigen quantification. For food antigen quantification, high-binding 96-well plates were coated with 100 µL of basolateral medium per well and incubated overnight at 4 °C. Food antigens were detected using antigen-specific antibodies against 19-kDa α-zein (MyBioSource), or Ara h 1 (MyBioSource). In competition experiments, organoid-derived monolayers were treated with 500 µg/mL CGM together with 50 µg/mL soluble FimH (MedChemExpress), FluoSpheres Microspheres at a 1:250 dilution, or 5 µg/mL anti-GP2 antibody (Proteintech).

**Particle Characterization Measurements**

Food antigens were prepared at a concentration of 5 mg/mL in Simulated Intestinal Fluid (SIF) and incubated overnight at 37 °C. Particle size and zeta potential were measured using a Zetasizer Nano ZS90 (Malvern Instruments).

**SEM Microscopy**

Samples were mounted to aluminum stubs using double sided carbon sticky tape then coated with a 20 nm layer of gold-palladium using a Leica EM SCD 500 sputter-coater. The samples were imaged in a Zeiss Sigma VP scanning electron microscope with a secondary electron detector. Images were collected at a resolution of 200 nm per pixel at an accelerating voltage of 3 kV.

**Untargeted Proteomics**

10 mg of bulk material was suspended in 200 uL of 70% methanol and shaken vigorously at 60 °C for 10 minutes. Samples were clarified by centrifugation at 5000 x g at 4C for 5 minutes. Proteins in the supernatant were then precipitated overnight at -20 C. On the following day, the pellets washed with methanol and redissolved in 4 M Guanidine Hydrochloride (GuHCl) with 10 mM tris(2-carboxyethyl)phosphine (TCEP) and 40 mM 2-Chloroacetamide (CAA) in 50 mM Triethylammonium bicarbonate (TEAB). The samples were heated to 95 C for 10 minutes in the dark to reduce and alkylate disulfide bonds. The GuHCl was diluted < 1M with 100 mM TEAB, and 10 mM Calcium Chloride (CaCl2) was added. The proteins were digested with chymotrypsin (1:30, Thermo Fisher) overnight at 28 °C. On the following day the digestion was stopped with 0.5% trifluoroacetic acid (TFA), and the peptides were desalted with Empore SDB-XC StageTips (CDS Analytical). Protein and peptide amounts were normalized with BCA and colorimetric

peptide assays (Pierce), respectively. After drying in a SpeedVac, the peptides were resuspended in 0.1% Formic acid (FA) in 5% Acetonitrile (ACN) and injected into a Thermo Fisher Easy nLC 1200 coupled to an Orbitrap Eclipse Tribrid mass spectrometer. Peptides were separated on a 50 cm uPAC UHPLC column (Thermo Fisher) at a flow rate of 300 nL/min. The mobile phases were 0.1% FA in water (A) and 0.1% FA in 80% ACN (B). The separation conditions were 4-30% B over 60 min followed by an increase to 50% B over 5 min. The column was then washed and re-equilibrated for the next sample. The mass spectrometer was operated in positive mode for data-dependent acquisition with a Nanospray Flex source for ionization. Full MS scans from 375-1500 m/z were acquired in the Orbitrap at 120k resolution. The most abundant precursors from each MS1 scan were subjected to HCD fragmentation (30 NCE), and the resulting MS2 scans were collected in the Orbitrap at 30k resolution over a total cycle time of 3 seconds. The isolation window was 1.0 m/z, and the dynamic exclusion time was 30 seconds. LC-MS data were analyzed with Byonic v5.10.62 (Protein Metrics) in ProteomeDiscoverer v3.2 (Thermo Fisher) against the Uniprot Zea mays reference proteome (Entry UP000007305_4577). Search settings included a precursor mass tolerance of 10 ppm and a fragment mass tolerance of 0.02 Da. Trypsin digestion was specified with a maximum of 3 missed cleavages and a minimum peptide length of 6. Modifications included fixed cysteine carbamidomethylation and variable methionine oxidation and N-terminal acetylation. Label-free quantitation based on precursor intensities was performed with the Minora feature finder and precursor quantification nodes in ProteomeDiscoverer, and protein abundances were normalized according to total peptide amount.

**Statistical analysis**

GraphPad Prism 11 was used for statistical analysis. Comparisons between two groups were performed using a t-test. Comparisons between multiple groups across one variable were analyzed using a one-factor ANOVA and comparisons with 2 variables were analyzed using a two-factor ANOVA. For ANOVA's with a significant P value, Tukey's multiple comparison testing was used for post-hoc analysis. $P < 0.05$ was considered statistically significant.

### Supplemental Figures

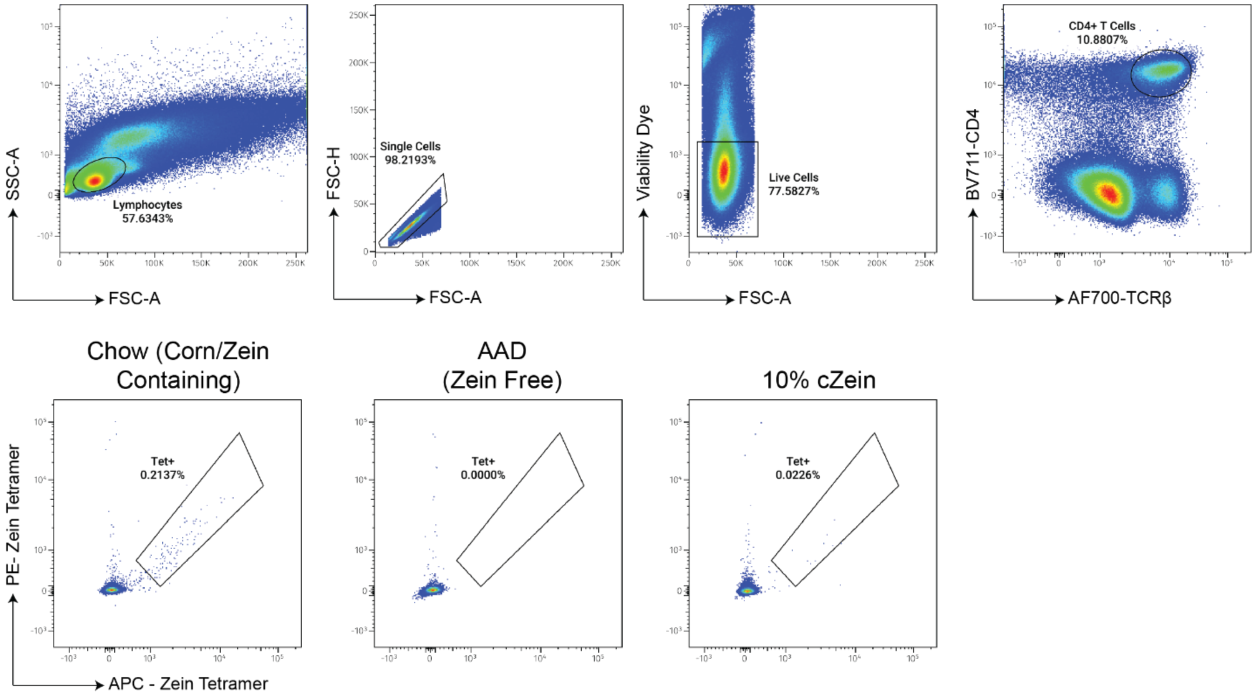


**Supplemental Figure 1: Representative flow gating.** Zein-specific CD4+ T cells were identified using flow cytometry analysis. AAD = Amino acid defined

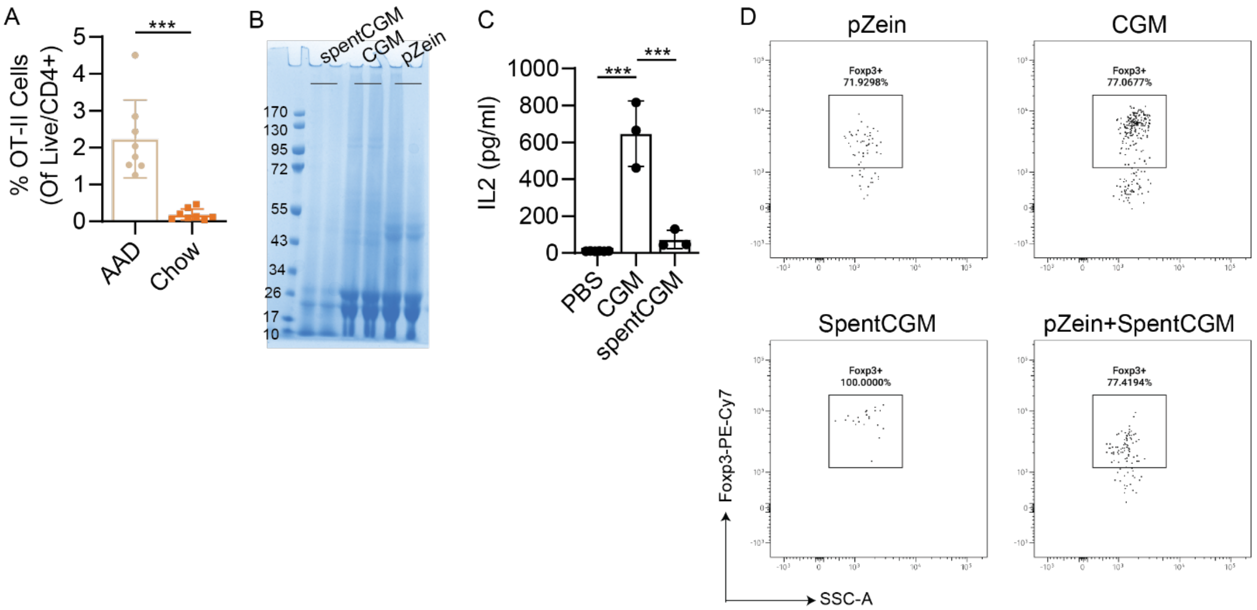


**Supplemental Figure 2: Antigen-specific T cell abundance and phenotype following feeding. (A)** Following adoptive transfer, OVA-specific T cells were more frequent in mice fed OVA on the background of AAD diet compared to chow diet (N=8/group). **(B)** spentCGM contains less zein, but is not fully zein deplete compared to CGM. **(C)** SpentCGM can minimally activate zein-reactive TCR transgenic lines, consistent with some residual zein protein in the spentCGM fraction. **(D)** Foxp3+ Treg is the dominant phenotype for all zein-specific T cells across dietary conditions. * P < 0.05, ** P < 0.01, *** P < 0.001. Error bars represent standard deviation.

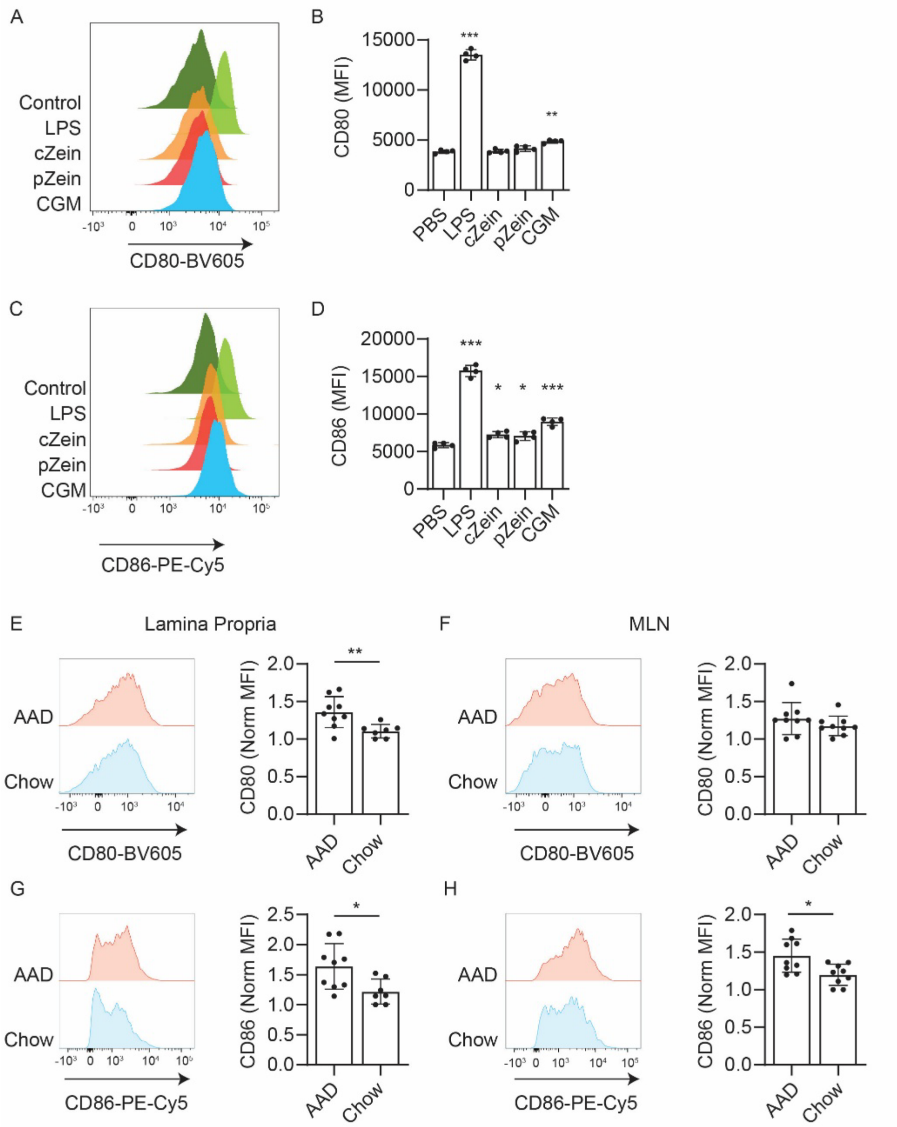


**Supplemental Figure 3: Lack of evidence for APC activation in response to zein. (A-D)** CD80 or CD86 levels in APCs following incubation with indicated stimuli *ex vivo* (N=4/group). **(E-H)** CD80 and CD86 levels in APCs from lamina propria or MLN *in vivo* (N=7-9/group). ** P < 0.01, *** P < 0.001. Error bars represent standard deviation.

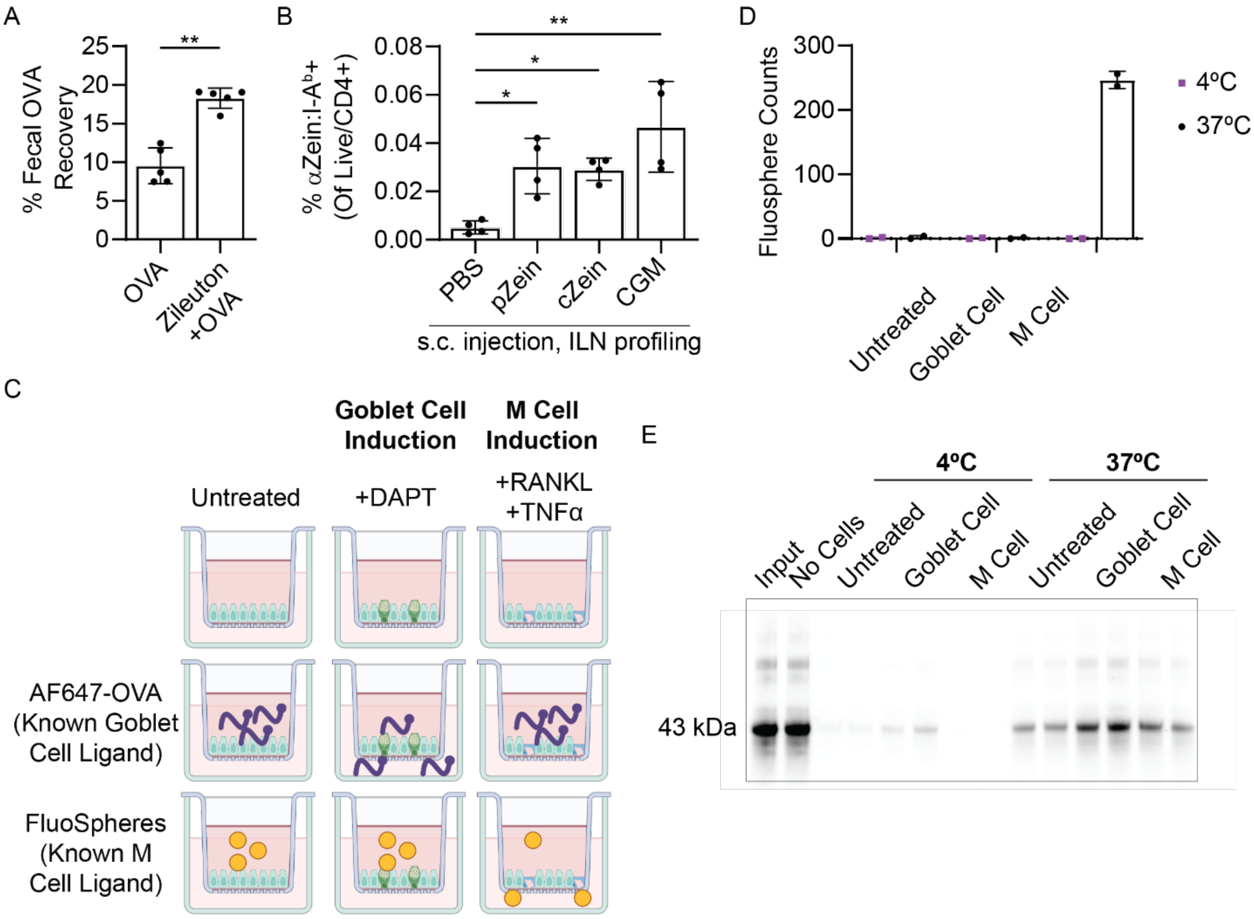


**Supplemental Figure 4: Validation that intestinal uptake limits Zein-specific T cell induction and organoid development. (A)** Mice were gavaged with OVA alone or zileuton+OVA. Fecal OVA recovery was measured using the FRAG assay (N=5/group). **(B)** Mice were injected with zein fractions emulsified in CFA and zein-specific T cells were measured in the inguinal lymph node (ILN) (N=4/group). **(C)** A transwell organoid assay was used to measure routes of intestinal antigen sampling. Induction with DAPT simulates goblet cell differentiation. Induction with RANKL and TNFα stimulates M cell differentiation. **(D)** Fluospheres, a known M cell ligand, are transported only in wells with induced M cells and only at 37°C (N=2/group). **(E)** AF647-OVA, a known goblet cell ligand, is transported only in wells with induced goblet cells at only at 37°C (N=2/group). * $P < 0.05$, ** $P < 0.01$, *** $P < 0.001$. Error bars represent standard deviation.

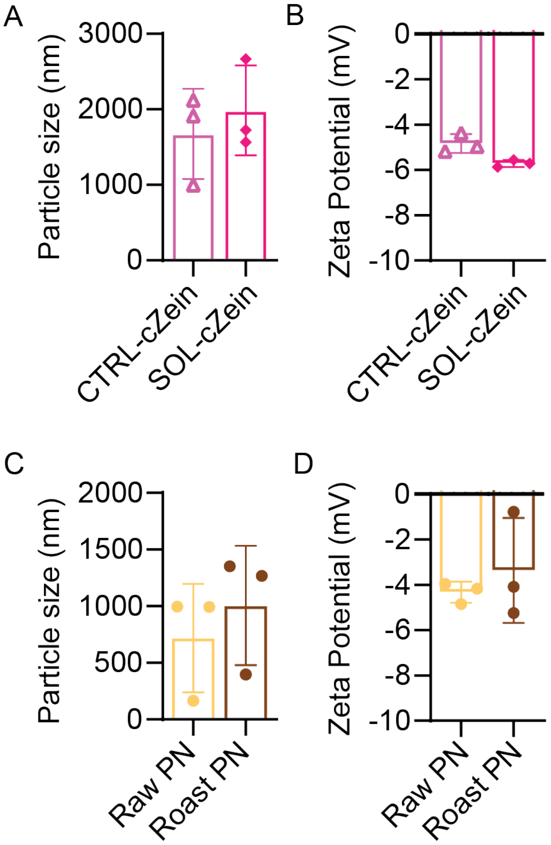


**Supplemental Figure 5: Particle analysis with solubilized zein and raw or roasted peanut.** **(A)** Particle size between CTRL-cZein and GLY-cZein (N=3/group). **(B)** Zeta potential between CTRL-cZein and GLY-cZein (N=3/group). **(C)** Particle size between raw and roast peanut (N=3/group). **(D)** Zeta potential between raw and roast peanut (N=3/group). * $P < 0.05$, ** $P < 0.01$, *** $P < 0.001$. Error bars represent standard deviation.
